# Nucleosomes and DNA methylation shape meiotic DSB frequency in Arabidopsis transposons and gene regulatory regions

**DOI:** 10.1101/160911

**Authors:** Kyuha Choi, Xiaohui Zhao, Christophe Lambing, Charles J. Underwood, Thomas J. Hardcastle, Heïdi Serra, Andrew J. Tock, Piotr A. Ziolkowski, Nataliya E. Yelina, Robert A. Martienssen, Ian R. Henderson

## Abstract

Meiotic recombination initiates via DNA double strand breaks (DSBs) generated by SPO11 topoisomerase-like complexes. Recombination frequency varies extensively along eukaryotic chromosomes, with hotspots controlled by chromatin and DNA sequence. To map meiotic DSBs throughout a plant genome, we purified and sequenced Arabidopsis SPO11-1-oligonucleotides. DSB hotspots occurred in gene promoters, terminators and introns, driven by AT-sequence richness, which excludes nucleosomes and allows SPO11-1 access. A strong positive relationship was observed between SPO11-1 DSBs and final crossover levels. Euchromatic marks promote recombination in fungi and mammals, and consistently we observe H3K4^me3^ enrichment in proximity to DSB hotspots at gene 5’-ends. Repetitive transposons are thought to be recombination-silenced during meiosis, in order to prevent non-allelic interactions and genome instability. Unexpectedly, we found strong DSB hotspots in nucleosome-depleted Helitron/Pogo/Tc1/Mariner DNA transposons, whereas retrotransposons were coldspots. Hotspot transposons are enriched within gene regulatory regions and in proximity to immunity genes, suggesting a role as recombination-enhancers. As transposon mobility in plant genomes is restricted by DNA methylation, we used the *met1* DNA methyltransferase mutant to investigate the role of heterochromatin on the DSB landscape. Epigenetic activation of transposon meiotic DSBs occurred in *met1* mutants, coincident with reduced nucleosome occupancy, gain of transcription and H3K4^me3^. Increased *met1* SPO11-1 DSBs occurred most strongly within centromeres and Gypsy and CACTA/EnSpm coldspot transposons. Together, our work reveals complex interactions between chromatin and meiotic DSBs within genes and transposons, with significance for the diversity and evolution of plant genomes.

## Introduction

Sexual eukaryotes reproduce via fusion of haploid gametes, which are produced by the specialized meiotic cell division. During meiosis a single round of DNA replication is coupled to two rounds of chromosome segregation. Additionally, during prophase of the first meiotic division, homologous chromosomes pair and recombine, which can result in reciprocal crossovers (Kauppi et al. 2004; Villeneuve and Hillers 2001). As a consequence of recombination and independent chromosome segregation, meiosis has a major effect on genetic variation within populations and the process of evolutionary adaptation (Barton and Charlesworth 1998; Hamilton 2002).

Meiotic recombination initiates via formation of DNA double strand breaks (DSBs), which can be repaired using a homologous chromosome to produce crossover or non-crossover products (Kauppi et al. 2004; Baudat et al. 2013; Villeneuve and Hillers 2001; Szostak et al. 1983). Meiotic DSBs are universally generated by SPO11 topoisomerase-like transesterases, which act as dimers to cleave opposite phosphodiester backbones using catalytic tyrosine residues (Neale et al. 2005; Keeney and Kleckner 1995; Pan et al. 2011; Keeney et al. 1997). In plants, SPO11-1 and SPO11-2 interact with MEIOTIC TOPOISOMERASE VIB (MTOPVIB), which forms a conserved catalytic core complex (Robert et al. 2016; Vrielynck et al. 2016; Hartung et al. 2007; Grelon et al. 2001). Following phosphodiester cleavage the SPO11 catalytic tyrosine remains covalently bound to the target site 5’-end (Neale et al. 2005; Keeney and Kleckner 1995; Pan et al. 2011). Endonucleases, including Sae2 and Mre11, then generate additional DNA backbone cuts 3’ to the DSB site that together with exonucleases, cause release of SPO11-oligonucleotide complexes (Garcia et al. 2011; Lam and Keeney 2014; Neale et al. 2005). Purification and sequencing of SPO11-oligonucleotides, which are typically ∼20–40 bases in length, has provided a high-resolution method to profile meiotic DSB patterns genome-wide in fungi and mammals (Pan et al. 2011; Lange et al. 2016; Fowler et al. 2014).

Meiotic DSB and crossover frequency vary extensively along eukaryotic chromosomes and typically concentrate in ∼1–2 kilobase hotspots (Baudat et al. 2013; Kauppi et al. 2004; Choi and Henderson 2015). Genetic and epigenetic information make varying contributions to control of hotspot locations and activities in different eukaryotic lineages (Baudat et al. 2013; Kauppi et al. 2004; Choi and Henderson 2015). For example, budding yeast DSB hotspots form predominantly in nucleosome-depleted regions in gene promoters and rarely in exons and terminators (Pan et al. 2011; Lam and Keeney 2015; Wu and Lichten 1994; Fan and Petes 1996). Local base composition, higher-order chromosome structure, transcription factor binding and ATM/ATR kinase signaling have further been shown to modify budding yeast DSB frequency (Lam and Keeney 2014; de Massy 2013; Cooper et al. 2016; Székvölgyi et al. 2015).

In contrast to budding yeast, mouse SPO11-oligonucleotides form at specific C-rich DNA sequence motifs that are bound by the meiotic protein PRDM9, which possesses a zinc finger array and a SET domain that catalyzes histone H3K4^me3^ and H3K36^me3^ (Lange et al. 2016; Mihola et al. 2009; Parvanov et al. 2010; Myers et al. 2010; Baudat et al. 2010; Grey et al. 2011; Brick et al. 2012; Grey et al. 2017; Powers et al. 2016). PRDM9-dependent SPO11 hotspots tend to show well-positioned flanking nucleosomes, which acquire H3K4^me3^ and H3K36^me3^ during meiosis (Grey et al. 2017; Baker et al. 2014; Lange et al. 2016; Brick et al. 2012; Powers et al. 2016). However, H3K4^me3^ levels do not correlate strongly with mouse or yeast SPO11-oligonucleotide levels, implying a downstream role for this chromatin mark (Tischfield and Keeney 2012; Lange et al. 2016). For example, budding yeast H3K4^me3^ is bound by the Spp1 COMPASS complex subunit, which simultaneously interacts with meiotic chromosome axis protein Mer3 and tethers chromatin loops to repair sites (Sommermeyer et al. 2013; Borde et al. 2009; Acquaviva et al. 2013). Furthermore, the mouse COMPASS subunit CXXC1 interacts with both PRDM9 and the IHO1 axis proteins, suggesting a conserved mechanism of chromatin loop-tethering during DSB repair (Imai et al. 2017).

Plant crossovers are enriched in euchromatin at the chromosome scale, and in proximity to gene promoters and terminators at the fine scale (Choi et al. 2013; Shilo et al. 2015; Drouaud et al. 2013; Wijnker et al. 2013; Horton et al. 2012; Hellsten et al. 2013; Fu et al. 2002; Choulet et al. 2014). Crossovers in plant genomes show positive associations with H3K4^me3^, histone variant H2A.Z (Choi et al. 2013; Wijnker et al. 2013; Shilo et al. 2015; Drouaud et al. 2013; Liu et al. 2009), A-rich and CTT/CNN-repeat DNA sequence motifs (Shilo et al. 2015; Choi et al. 2013; Wijnker et al. 2013), and can be directly suppressed by acquisition of heterochromatic modifications, such as DNA methylation and H3K9^me2^ (Yelina et al. 2015). However, genome-wide meiotic DSB patterns and their relation to chromatin, DNA sequence and crossover frequency have yet to be reported in a plant genome.

Despite deep conservation of core meiotic factors, such as SPO11, many aspects of genome architecture, chromatin and recombination vary between eukaryotes. For example, budding yeast possesses point centromeres, whereas large, regional centromeres surrounded by repetitive heterochromatin are more common in other eukaryotes (Bloom 2014; Copenhaver et al. 1999; Vincenten et al. 2015; Malik and Henikoff 2009). Equally, although transposable elements are ubiquitous, their diversity and abundance varies between species (Feschotte and Pritham 2007; Beauregard et al. 2008; McClintock 1956). Transposons are typically heterochromatic and show RNA polymerase-II suppression, caused by epigenetic modifications (Slotkin and Martienssen 2007). Repetitive sequences are also frequently crossover-suppressed during meiosis, in order to limit non-allelic homologous recombination and genome instability (Sasaki et al. 2010). However, evidence exists for specific transposon families promoting meiotic recombination in plants, fungi and animals (Myers et al. 2005; Shi et al. 2010; Sasaki et al. 2013; Horton et al. 2012; Yandeau-Nelson et al. 2005). For example, meiotic gene conversion, although not crossovers, has been observed in maize centromeric transposons (Shi et al. 2010), which indicates DSB formation and interhomolog repair. However, the extent to which plant transposons and repetitive sequences initiate meiotic recombination genome-wide has remained unclear.

To further explore relationships between recombination and chromatin, in genes versus repeats, we mapped meiotic DSBs and crossovers throughout the ∼135 megabase *Arabidopsis thaliana* genome, which contains diverse DNA and RNA transposons (Supplemental Fig. S1) (Buisine et al. 2008; Quadrana et al. 2016; Stuart et al. 2016; Kawakatsu et al. 2016; Slotkin and Martienssen 2007). We show that Arabidopsis meiotic DSB hotspots are concentrated in gene promoters, terminators and introns. We also observe strong DSB hotspots inside specific families of DNA transposons, which are enriched within gene regulatory sequences. We show that nucleosome occupancy, driven by AT-sequence richness, is a major determinant of DSB hotspot strength and location in both genes and repeated sequences. Using the *met1* DNA methylation mutant, we demonstrate coordinate epigenetic remodeling of transcription, chromatin and recombination. Activation of meiotic DSBs in *met1* occurs most strongly in centromeric heterochromatin and specific Gypsy and EnSpm/CACTA transposon families. Together, our work reveals both conserved and plant-specific aspects to the meiotic DSB landscape and its relationship to chromatin.

## Results

### Purification and sequencing of Arabidopsis SPO11-1-oligonucleotides

In order to map meiotic DSBs throughout the Arabidopsis genome we sought to purify and sequence SPO11-1-oligonucleotides (Pan et al. 2011; Grelon et al. 2001). We generated a 6×Myc translational fusion at the C-terminus of Arabidopsis *SPO11-1*, driven by the endogenous promoter, which fully complements *spo11-1* fertility and crossover frequency, measured using fluorescent recombination reporter lines (Fig. 1A–1B and Supplemental Table S1) (Berchowitz and Copenhaver 2008; Grelon et al. 2001). To analyze SPO11-1-Myc during meiosis we performed immunocytology using α-Myc antibodies. SPO11-1-Myc foci were detected from leptotene until pachytene stage, associated with the meiotic chromosome axis, which was visualized by co-immunostaining for the ASYNAPTIC1 (ASY1) HORMA domain protein (Fig. 1C and Supplemental Fig. S2). SPO11-1-Myc foci showed a comparable number (mean=204.6 foci, *n=*10) and duration to those reported for its binding partner MTOPVIB (Fig. 1C and Supplemental Fig. S2) (Vrielynck et al. 2016). No α-Myc signal was detected above background in wild type meiotic cells, or in *SPO11-1-Myc* somatic cells (Fig. 1C). Therefore, SPO11-1-Myc is functional and accumulates on meiotic chromosomes, coincident with endogenous DSB formation (Vrielynck et al. 2016; Sanchez-Moran et al. 2007).

**Figure 1.**
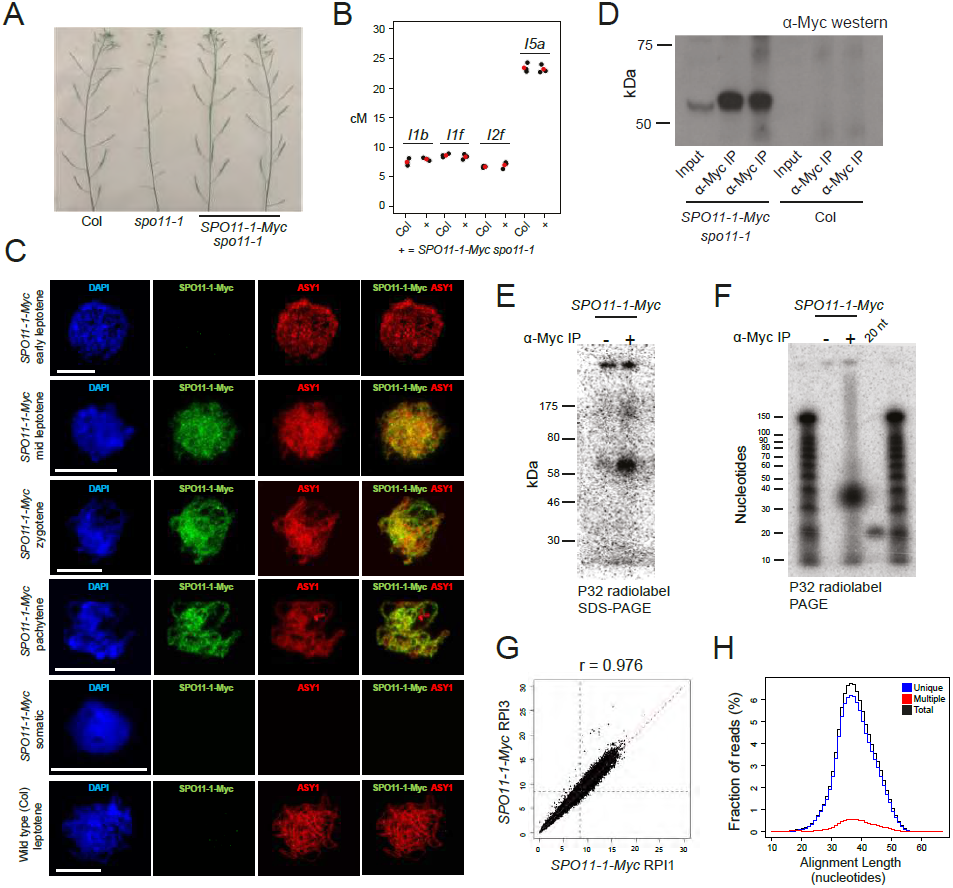
Purification and sequencing of Arabidopsis SPO11-1-oligonucleotides. (A) Inflorescences from wild type (Col), *spo11-1* and *SPO11-1-Myc spo11-1* plants. (B) Crossover frequency (cM) measured using fluorescent crossover reporter lines in Col or *SPO11-1-Myc spo11-1*, with mean values in red. (C) Nuclei from *SPO11-1-Myc* or Col pollen mother cells immunostained for α-Myc (green) or α-ASY1 (red), and stained for DAPI (blue). Scale bars=10μM. (D) α-Myc western blotting from *SPO11-1-Myc* or Col extracts, before and after α-Myc immunoprecipitation (α-Myc-IP). (E) Detection of end-radiolabelled SPO11-1-Myc complexes following immunoprecipitation and SDS polyacrylamide gel electrophoresis (SDS-PAGE). (F) Detection of purified SPO11-1-oligonucleotides following proteinase K digestion of immunoprecipitates and polyacrylamide gel electrophoresis (PAGE). A labeled 20 base oligonucleotide (20 nt) was run alongside as a size control. (G) Correlation of SPO11-1 in adjacent 10 kb windows for wild type libraries RPI1 and RPI3 (Supplementary Table 2). Blue dotted lines indicate genome average values. The Pearson’s correlation coefficient (*r*) is printed above. (H) Histogram showing lengths of uniquely aligning (blue), multiple-aligning (red) and total (black) SPO11-1 reads.

Following protein extraction from meiotic-stage floral buds, SPO11-1-Myc was detectable as a ∼54 kDa band using western blotting (Fig. 1D). Oligonucleotides covalently attached to SPO11-1-Myc can be radioactively 3’-end labeled using terminal transferase (Neale and Keeney 2009), which revealed 60–70 kDa complexes (Fig. 1E). No signal was observed when the protocol was repeated without antibody (Fig. 1E). Following proteinase K digestion of SPO11-1-Myc immunoprecipitates and PAGE separation, we detected radiolabelled SPO11-1-oligonucleotides ∼35–45 bases in length (Fig. 1F). SPO11-1-oligonucleotides were gel purified and used to generate sequencing libraries, using a protocol adapted from budding yeast (Supplemental Fig. S3) (Pan et al. 2011). Three biological replicate wild type (*SPO11-1-Myc spo11-1*) libraries were sequenced to high depth (11–28 million mapped reads), which showed significant correlation (Fig. 1G, Supplemental Fig. S4 and Supplemental Tables S2–S3). For example, Pearson’s *r* between replicates was 0.97–0.98 at the 10 kb scale (Supplemental Table S3). The majority (92.2–93.4%) of SPO11-1-oligonucleotide reads (hereafter called SPO11-1) aligned uniquely, and multiple-mapped reads with equal alignment scores were randomly assigned (Fig. 1H and Supplemental Table S2).

### Genomic landscapes of SPO11-1-oligonucleotides, crossovers, euchromatin and heterochromatin

We analyzed SPO11-1 levels in 10 kb windows and plotted DSB frequency throughout the Arabidopsis genome (Fig. 2A–2C). Consistent with broad-scale patterns of crossover recombination (Choi et al. 2013; Giraut et al. 2011; Salomé et al. 2012), SPO11-1 is highest in the euchromatic chromosome arms, lowest in the centromeres (Fig. 2A), and shows a positive correlation with genes (*r*=0.777) and a negative correlation with transposon density (*r*=-0.816). To compare with epigenetic marks, we performed ChIP-seq for the gene-enriched histone modification H3K4^me3^, which was positively correlated with SPO11-1 (*r*=0.700), whereas centromere-enriched DNA methylation was negatively correlated (*r*=-0.831) (Fig. 2A–2B and Supplemental Table S6) (Yelina et al. 2015). This is consistent with chromatin playing a major role in shaping the Arabidopsis DSB landscape, at the chromosome scale.

**Figure 2.**
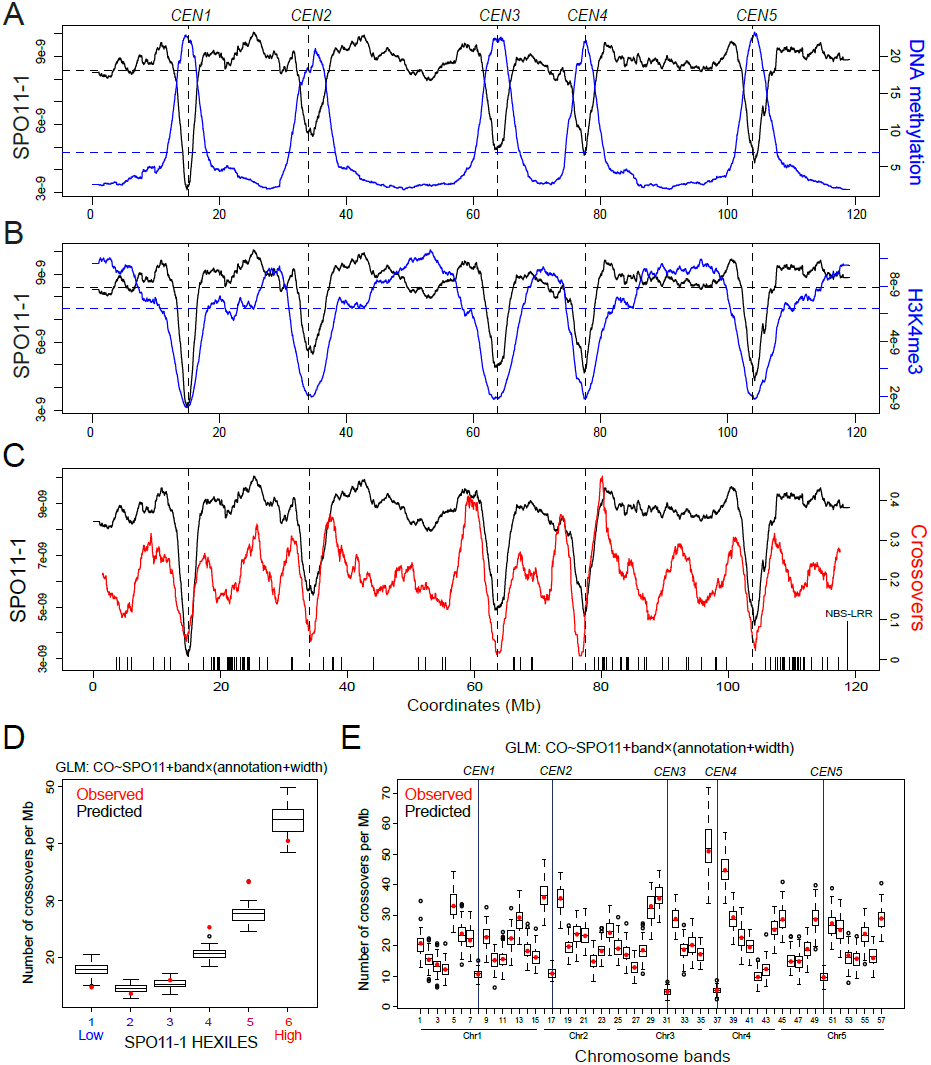
Genomic landscape of SPO11-1 DSBs, crossovers, euchromatin and heterochromatin. (A) SPO11-1 (black) and DNA methylation (blue) density throughout the Arabidopsis genome, with centromeres indicated by vertical dotted lines. Horizontal dotted lines represent mean values. (B) As for (A), but plotting SPO11-1 (black) and H3K4^me3^ (blue). (C) As for (A), but plotting SPO11-1 (black) and crossover frequency (red). Crossovers were identified using genotyping-by-sequencing of Col×Ler F_2_ plants. X-axis ticks indicate the positions of NBS-LRR gene homologs. (D) Observed (red dots) crossover overlap per megabase for SNP intervals, grouped according to SPO11-1 hexiles (1=low SPO11-1, 6=high SPO11-1). Boxplots show the range of predicted crossover overlap values based on the generalized linear model (GLM) formula: CO∼SPO11+band×(annotation+width). (E) As for (D), but showing observed and predicted crossover overlaps per megabase, according to two megabase chromosomal bands.

In order to compare meiotic DSB levels with the frequency of final crossover products, we used 2,499 crossover events mapped in 363 Col×Ler F_2_ plants by genotyping-by-sequencing(Fig. 2C–2E and Supplemental Table S4) (Choi et al. 2016; Rowan et al. 2015). Crossovers were mapped between Col/Ler SNPs to a mean resolution of 970 bp. At the chromosome scale there was a positive correlation between SPO11-1 and crossover frequency (*r*=0.593) (Fig. 2C). However, there was also significant variation in the ratio of SPO11-1 to crossovers along the chromosome arms (Fig. 2C), which may reflect modification of recombination downstream of DSB formation, for example by polymorphism, or additional features of chromosome architecture. We used a logistic model to analyze the likelihood of observing crossovers relative to other genome features. This revealed a strong positive effect for SPO11-1 levels (10.87, *P=*9.36×10^-87^), with weaker but significant effects for chromosome position and sequence annotation (Fig. 2D–2E and Supplemental Table S5). Therefore, overall higher levels of initiating meiotic DSBs associate with higher final crossover levels.

### SPO11-1 DSB hotspots in nucleosome-depleted gene regulatory regions

Budding yeast meiotic DSB hotspots occur in nucleosome-depleted regions within gene promoters (Pan et al. 2011; Lam and Keeney 2015; Wu and Lichten 1994; Fan and Petes 1996; Nicolas et al. 1989), whereas mammalian PRDM9-dependent DSB hotspots tend to be located intergenically at specific C-rich sequence motifs (Myers et al. 2008; Brick et al. 2012; Kong et al. 2010; Lange et al. 2016). Therefore, we analyzed Arabidopsis SPO11-1-oligonucleotides in relation to gene transcriptional start sites (TSSs) and termination sites (TTS) (Fig. 3A–3B). We also analyzed nucleosome occupancy by performing micrococcal nuclease digestion of chromatin, followed by sequencing of mononucleosomal DNA (MNase-seq) (Fig. 3A–3B and Supplemental Table S7) (Choi et al. 2016). Similar to budding yeast, SPO11-1 was highest in Arabidopsis nucleosome-free regions located in gene promoters (Fig. 3A–3C). Interestingly we also observe strong DSB hotspots in nucleosome-free terminators, where plant crossover hotspots are also observed (Fig. 3A–3C) (Choi et al. 2013; Wijnker et al. 2013). A further difference is that Arabidopsis genes possess on average 6.7 exons (Cheng et al. 2017), whereas budding yeast genes lack introns (Pan et al. 2011). We observe that Arabidopsis introns have higher SPO11-1 and lower nucleosomes compared with exons (Supplemental Fig. S5B–S5D). However, SPO11-1 is overall suppressed within relatively nucleosome-occupied gene bodies, compared with flanking nucleosome-depleted promoter and terminator regions (Fig. 3A–3C and Supplemental Fig. S5A). As expected, H3K4^me3^ shows prominent enrichment at the +1 nucleosome position, immediately downstream of TSS, within gene bodies (Fig. 3B) (Zhang et al. 2009).

**Figure 3.**
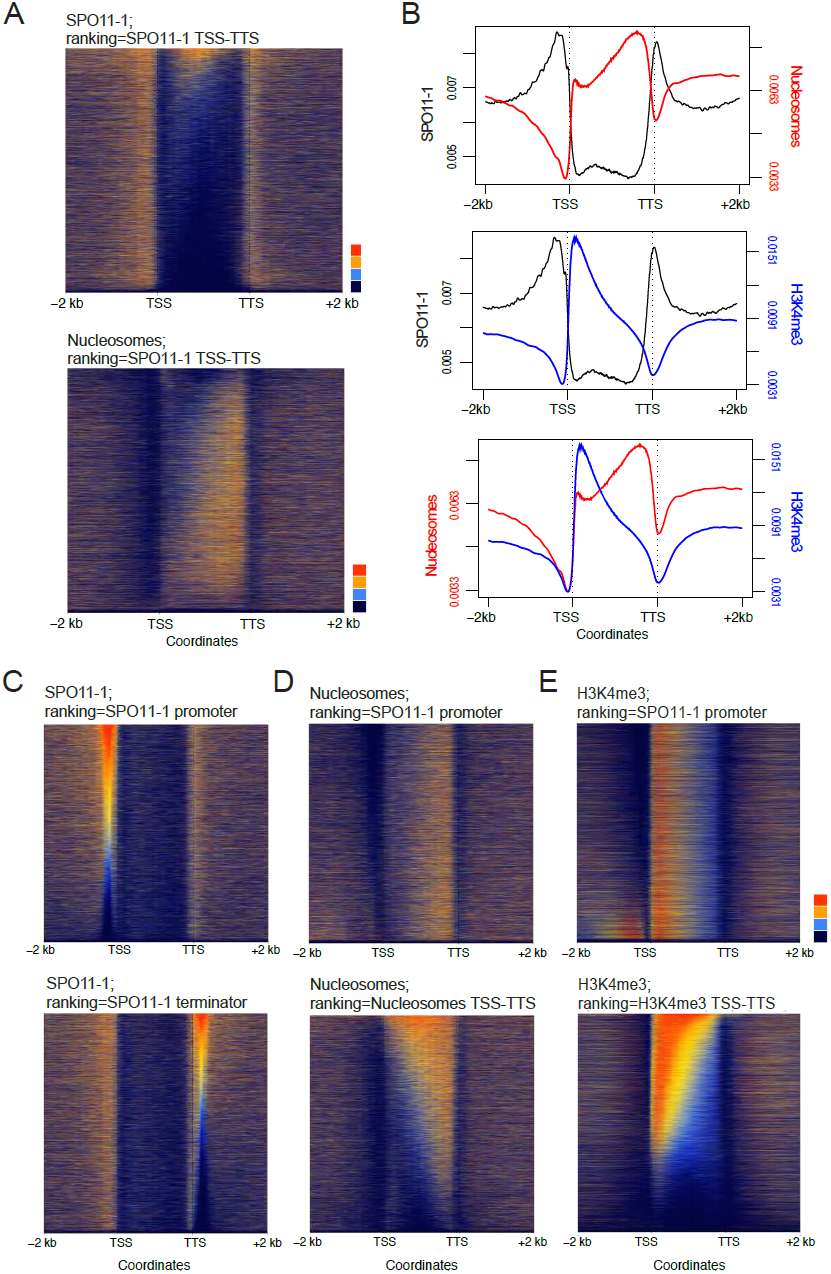
SPO11-1 DSB hotspots in gene promoter and terminator nucleosome-free regions. (A) Heat maps of SPO11-1-oligonucleotides (upper) and nucleosomes (lower) within gene transcriptional units (between transcriptional start (TSS) and termination (TTS) sites) and 2 kb flanking regions. Each row represents an individual gene, which have been ordered by SPO11-1-oligonucleotide normalized coverage values between TSS and TTS. SPO11-1 and nucleosome values equal to defined quantiles were mapped linearly to a vector of six colors (dark blue (lowest), blue, light blue, yellow, orange, red (highest)). (B) Density of SPO11-1-oligonucleotides (black), nucleosome occupancy (MNase-seq, red), or H3K4^me3^ (ChIP-seq, blue) in wild type, across gene transcriptional units (TSS to TTS) and in flanking 2 kb windows. (C) Heat maps as for (A), but showing SPO11-1 ranked by SPO11-1 levels in gene promoters (upper, −500 bp upstream of TSS) or gene terminators (lower, +500 bp downstream of TTS). (D) Heat maps as for (A), but showing nucleosomes ranked by SPO11-1 levels in gene promoters (upper) or by nucleosomes within TSS–TTS (lower). (E) Heat maps as for (A), but showing H3K4^me3^ ranked by SPO11-1 levels in gene promoters (upper) or by H3K4^me3^ within TSS–TTS (lower).

To investigate control of DSB levels we ranked genes according to SPO11-1 in 500 bp windows upstream of gene TSS (promoters), or downstream of TTS (terminators) (Fig. 3C). Levels of promoter SPO11-1 did not strongly associate with terminator levels, showing that meiotic DSBs vary independently at opposite ends of genes (Fig. 3C). We used the SPO11-1 promoter ranking to look at associated variation in nucleosome occupancy (MNase) and H3K4^me3^ levels. High SPO11-1 promoters strongly associate with lower promoter nucleosome occupancy, consistent with DNA accessibility being a major determinant of Arabidopsis DSB levels (Fig. 3D). In contrast, H3K4^me3^ levels within genes did not show a strong association with promoter SPO11-1 levels (Fig. 3E). This supports a recombination-promoting role for H3K4^me3^ downstream of DSB formation, consistent with analysis of mouse and budding yeast SPO11-oligonucleotides (Tischfield and Keeney 2012; Lange et al. 2016).

### SPO11-1 hotspots and coldspots in transposable elements

To explore meiotic DSB levels within repetitive sequences we selected 29,150 transposable elements from 10 DNA and RNA families for analysis (Supplemental Fig. S1 and Supplemental Table S8) (Buisine et al. 2008). Extensive SPO11-1 variation was observed between transposon families, with high DSB levels in Helitrons, which transpose via rolling-circle replication, and Pogo/Tc1/Mariner and MuDR ‘cut-and-paste’ DNA transposons (Fig. 4A and Supplemental Table S8) (Kapitonov and Jurka 2001; Slotkin and Martienssen 2007). In contrast, retrotransposons that replicate via RNA intermediates, including LTR and non-LTR families, were SPO11-1 coldspots (Fig. 4A and Supplemental Table S8) (Beauregard et al. 2008). As observed for genes (Fig. 3), variation in transposon family SPO11-1 negatively correlated with nucleosome occupancy (*r=*-0.96) (Fig. 4B and Supplemental Table S8). We divided transposons into six groups (hexiles) after ranking by within element SPO11-1 levels (Figure 4C; hexile 1=highest, hexile 6=lowest). This grouping showed strong correlations between higher SPO11-1 and reduced transposon lengths (*r*=-0.80), lower nucleosome occupancy (*r*=-0.94), greater DNA (*r*=0.95) and fewer RNA transposons (*r*=-0.95) (Fig. 4C and Supplemental Tables S8-S9). At the chromosome scale, high SPO11-1 transposons (e.g. Helitrons and Pogo/Tc1/Mariner) show elevated density in the chromosome arms and pericentromeres, whereas low SPO11-1 transposons (e.g. Gypsy LTR) are centromere-enriched (Fig. 4D). Differences in DSB activity between transposon families are also evident locally, for example comparing a nucleosome-dense retroelement coldspot *ATCOPIA4* with an adjacent cluster of nucleosome-depleted Helitron hotspots (Fig. 4E–4F). Many DSB hotspot DNA transposons are short, non-autonomous fragments, although high SPO11-1 was also observed within full length Helitron and *Lemi1* Pogo transposons (Supplemental Fig. S6A–S6D) (Feschotte and Mouchès 2000; Kapitonov and Jurka 2001). Hence, despite the expectation that transposons would be suppressed for meiotic DSBs, we observe that specific families of repetitive elements are nucleosome-depleted SPO11-1 hotspots.

**Figure 4.**
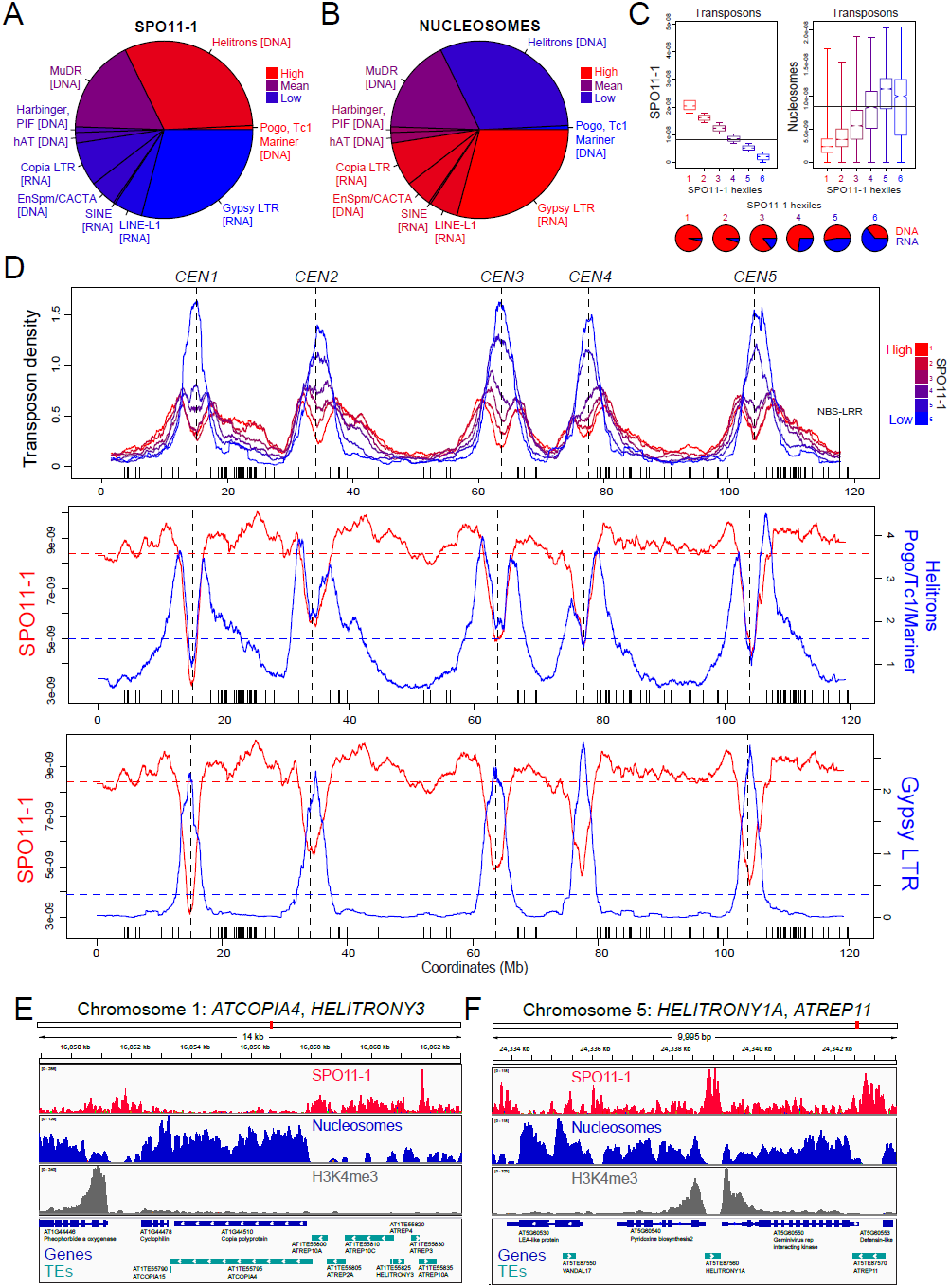
Meiotic recombination and chromatin variation between Arabidopsis transposons. (A) Pie chart showing Arabidopsis transposon families, with slice size proportional to physical length, and color-coded according to SPO11-1 levels. The color equivalent to the genome-wide mean value is inset. (B) As for (A), but showing nucleosome occupancy (MNase-seq). (C) Box plots showing SPO11-1 and nucleosome occupancy, according to transposon SPO11-1 hexile groups, with horizontal lines indicating the genome average value. Inset pie charts show the proportion of DNA (red) and RNA (blue) transposons for each SPO11-1 hexile. (D) Density of transposons through the Arabidopsis genome according to SPO11-1 hexile (red=highest SPO11-1, blue=lowest SPO11-1). X-axis ticks indicate NBS-LRR gene homologs. Plotted beneath are SPO11-1 (red) versus Helitron/Pogo/Tc1/Mariner class DNA transposons (blue), or Gypsy RNA transposons (blue). (E)–(F) Close-ups of chromosomal regions showing SPO11-1 (red), nucleosomes (blue) and H3K4^me3^ (grey) density, relative to gene (dark blue) and transposon (light blue) annotation shown beneath. Note in (F), the presence of a *DEFENSIN* gene At5g60553 associated with a Helitron *ATREP11* hotspot.

### Nucleosomes, DNA sequence and SPO11-1 within genes and transposons

To further investigate spatial relationships between meiotic DSBs, chromatin and DNA sequence, we analyzed 4 kb windows around gene TSS and TTS, or transposon start and end coordinates, each according to SPO11-1 hexile groups (Fig. 5A–5B and Tables S9– S11). Again, a strong negative relationship between SPO11-1 and nucleosome occupancy was observed in both genes and transposons (Fig. 5A–5B and Supplemental Tables S9– S11). In the high SPO11-1 regions, we also observe quantitative enrichment of AT-rich sequence motifs that have previously been associated with high crossovers (Fig. 5A–5B and Supplemental Tables S6–S8) (Horton et al. 2012; Choi et al. 2013; Shilo et al. 2015; Wijnker et al. 2013). As AT-sequence richness is known to exclude nucleosomes (Segal and Widom 2009), we propose that these motifs cause higher SPO11-1 accessibility via this effect, leading to higher DSB formation and crossover frequency (Fig. 5A–5B).

**Figure 5.**
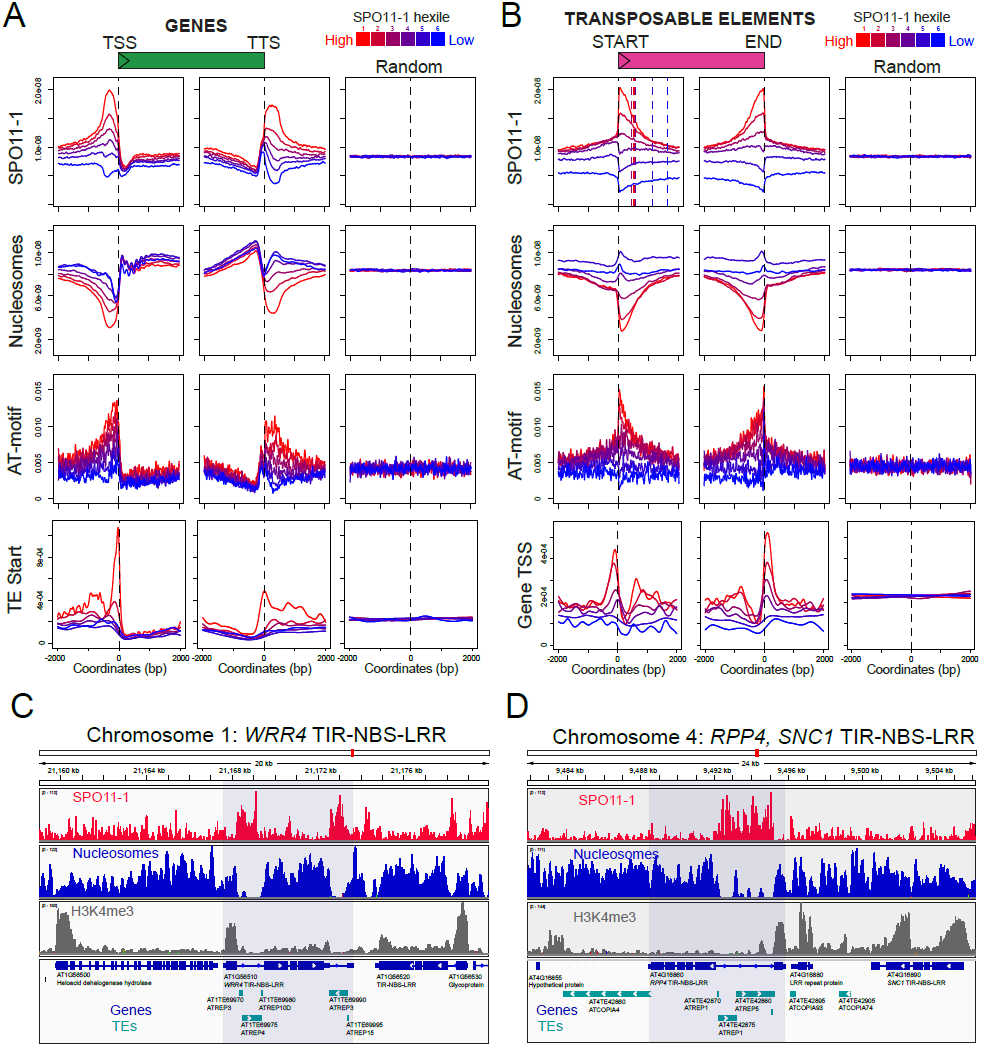
Nucleosomes, AT-sequence motifs and SPO11-1 DSBs within genes and transposons. (A) Density of SPO11-1, nucleosomes, AT-motifs (Choi et al. 2013), and TE start coordinates in 4 kb windows around gene transcriptional start sites (TSSs) or termination sites (TTSs), or the same number of random (Random) positions. Genes are grouped according to SPO11-1 promoter or terminator hexiles (red=highest, blue=lowest).(B) As for (A) but analyzing transposon SPO11-1 hexiles, and showing gene TSS proximity.(C) Close-ups of chromosomal regions showing SPO11-1 (red), nucleosomes (blue) and H3K4^me3^ (grey), relative to gene (dark blue) and transposon (light blue) annotations. The *WRR4* TIR-NBS-LRR resistance gene is highlighted which contains transposon hotspots within its introns. (D) As for (C), with the *RPP4* TIR-NBS-LRR resistance gene highlighted, which contains intronic hotspot transposons.

We also note that high SPO11-1 genes and transposons show close proximity to one another (Fig. 5A–5B). Helitron transposons are known to insert into AT dinucleotides (Kapitonov and Jurka 2001), and *Lemi1* Pogo transposons insert into TA dinucleotides (Guermonprez et al. 2008). Therefore, transposon integration site preference likely contributes to DNA element enrichment in AT-rich gene promoters and terminators, where they further contribute to nucleosome exclusion and high meiotic DSB levels (Fig. 5A–5B). High recombination rates may also provide an explanation for the tendency of DSB hotspot transposons to be shorter (Supplemental Tables S8–S9), due to promotion of non-homologous recombination and sequence rearrangement (Sasaki et al. 2010). Together, these findings reveal intimate connections between DNA sequence, chromatin and recombination around Arabidopsis genes and transposons.

### Meiotic DSB hotspot transposons are enriched in proximity to immunity genes

To investigate genes associated with high DSB levels, we tested for enrichment of Gene Ontology (GO) terms, following ranking by promoter SPO11-1 levels (Fig. 3C). This revealed a strong association with biotic defense GO terms (Supplemental Table S12), which was driven by high recombination *DEFENSIN* genes (Supplemental Fig. S6E–S6F). *DEFENSINS* encode small cysteine-rich peptides with roles in antimicrobial defense and pollen-pistil interactions (Silverstein et al. 2005). Further association of recombination hotspots and immunity genes is evident at the chromosome scale, where high SPO11-1 transposons show elevated density within the nucleotide binding site-leucine rich repeat (NBS-LRR) immune gene clusters on the right arms of chromosomes 1 and 5 (Fig. 4D) (Choi et al. 2016), and 73 of 197 NBS-LRR genes are within 500 base pairs of DSB hotspot transposons (Supplemental Table S13). For example, the NBS-LRR crossover hotspots *RAC1* and *HRG1* are flanked by Helitron and MuDR hotspot transposons, respectively (Supplemental Fig. S6G–S6H and Supplemental Table S13) (Choi et al. 2016). Further examples include the *RPP4* and *WRR4* oomycete resistance genes, which contain strong *ATREP* Helitron DSB hotspots within their introns (Fig. 5C–5D and Supplemental Table S13) (van der Biezen et al. 2002; Borhan et al. 2008). As Arabidopsis NBS-LRR genes are sites of natural structural diversity and DNA methylation polymorphism in populations (Kawakatsu et al. 2016; Quadrana et al. 2016; Stuart et al. 2016), we propose that gene-proximal DNA transposons may act as meiotic recombination enhancers, contributing to the high levels of genetic and epigenetic variation observed at these loci.

### Epigenetic remodeling of SPO11-1 DSBs, chromatin and transcription in *met1* DNA methylation mutants

Heterochromatic marks, such as DNA methylation, play critical roles in transcriptionally silencing transposable elements and thereby limiting their proliferation within eukaryotic genomes (Slotkin and Martienssen 2007). To directly investigate the role of heterochromatin on transposon recombination, chromatin and transcription, we compared SPO11-1, nucleosomes, H3K4^me3^ and RNA expression genome-wide in wild type and *met1*. *MET1* encodes the major CG sequence context maintenance DNA methyltransferase in Arabidopsis (Stroud et al. 2013; Saze et al. 2003; Kankel et al. 2003). In *met1* mutants cytological decondensation of heterochromatin occurs, together with elevated transposon transcription and mobility (Mathieu et al. 2007; Saze et al. 2003; Kato et al. 2003). We therefore sought to test whether related changes in heterochromatic meiotic DSBs occur in *met1*. For all experiments we used the null *met1-3* allele, which was isolated in a Columbia background (Saze et al. 2003).

At the chromosome-scale *met1-3* shows pronounced loss of CG DNA methylation within the centromeric regions (Stroud et al. 2013). We observe that this is mirrored by an increased centromeric SPO11-1 differential between *met1* and wild type (ΔSPO11-1) (Fig. 6A). The *met1* ΔSPO11-1 differential also strongly negatively correlates with the *met1* nucleosome differential (*r*=-0.879), and positively with the *met1* H3K4^me3^ differential (*P*=0.837) (Fig. 6A). This shows that loss of CG DNA methylation causes broad-scale gain of both meiotic SPO11-1 DSBs and euchromatic chromatin states (reduced nucleosome occupancy and increased H3K4^me3^) within the *met1* centromeric regions. Regions showing high *met1* ΔSPO11-1 differential also strongly correlate with the densities of Gypsy (*r=*0.913) and EnSpm/CACTA (*r*=0.892) transposons, which are SPO11-1 coldspots in wild type (Figs. 4A and 6A).

**Figure 6.**
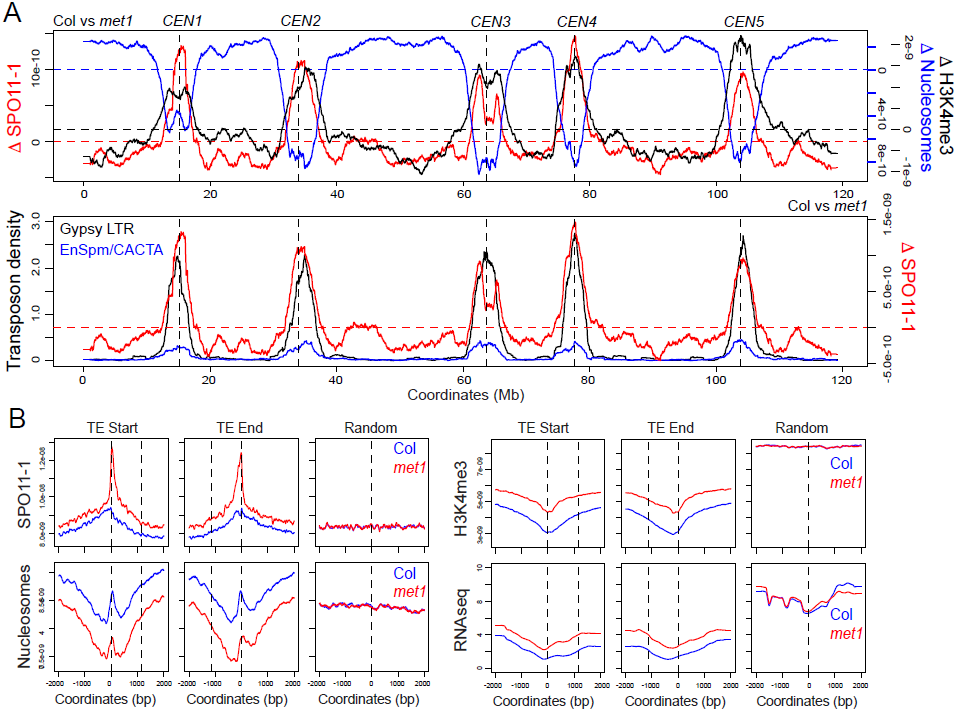
Coordinate epigenetic remodeling of chromatin, transcription and meiotic DSBs in *met1* DNA methylation mutants. (A) Differential (Δ) signal of SPO11-1 (red), nucleosomes (blue) and H3K4^me3^ (black) in *met1* compared with wild type (Col), throughout the Arabidopsis genome. Horizontal dotted lines indicate zero differential. Centromeres are indicated by vertical dotted lines. The lower plot shows ΔSPO11-1 (red) compared with Gypsy (black) and EnSpm/CACTA (blue) transposon densities. (B) SPO11-1, nucleosomes, H3K4^me3^ or RNAseq data in Col (blue) versus *met1* (red), analyzed in 4 kb windows around the start and end of those transposons with positive ΔSPO11-1 values (n=12,224), or the same number of random positions. The mean width of TEs analyzed is indicated by the vertical dotted lines.

To analyze changes in recombination at the fine-scale, we compared SPO11-1 levels within transposons between wild type and *met1* (Fig. 6B). 12,224 transposons (41.9%) showed net gain of SPO11-1 in *met1* (Fig. 6B). These recombination-activated transposons also show significantly reduced nucleosome occupancy, elevated H3K4^me3^ and increased transcription in *met1* (ANOVA all *P=*<2.50×10^-6^) (Fig. 6B). This is consistent with the trends observed at chromosome scale, and demonstrate that loss of CG methylation causes transposons to gain euchromatic features and increase meiotic recombination initiation. These trends are also evident at specific transposable elements. For example, the *ATENSPM9* and *ATENSPM10* EnSpm/CACTA and *ATGP3* Gypsy transposons show coordinate activation of transcription [fig6] and meiotic DSBs in *met1*, in addition to showing reduced nucleosome occupancy and gain of H3K4^me3^ (Fig. 7A).

**Figure 7.**
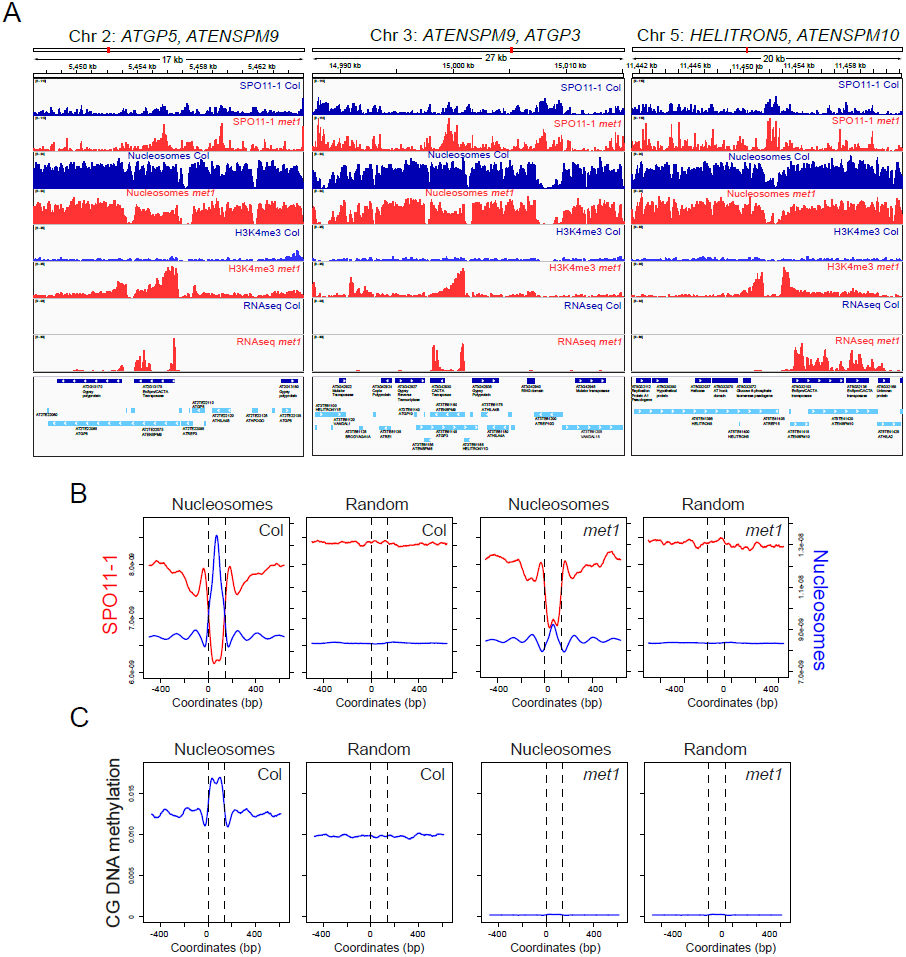
Fine-scale epigenetic remodeling of *met1* transposon chromatin, transcription and recombination. (A) Close-up of chromosomal regions showing SPO11-1, nucleosomes, H3K4^me3^ and RNAseq data, relative to gene (dark blue) and transposon (light blue) annotation, for Col (blue) and *met1* (red). (B) Plots analyzing SPO11-1 (red) and nucleosomes (blue) in Col and *met1* for highly positioned nucleosomes that are differentially occupied in *met1* (n=30,267), or the same number of random positions. (C) As for (B), but analyzing CG DNA methylation (blue) in wild type and *met1*.

To further analyze the interaction of chromatin structure and meiotic DSBs, we identified 74,401 highly positioned nucleosomes in wild type from our MNase-seq data. Of these positions 30,276 showed reduced nucleosome occupancy in *met1* (Fig. 7B). These nucleosome positions show high CG methylation in wild type, with peak ^m^CG levels immediately flanking the central nucleosomal peak (Fig. 7C). In *met1*, CG methylation is lost at these positions, which is coincident with significantly reduced nucleosome occupancy and increased SPO11-1 DSBs (ANOVA all *P=*<2.2×10^-16^) (Fig. 7B). Taken together, this demonstrates coordinate remodeling of chromatin, histone modifications, transcription and meiotic recombination, caused by loss of CG DNA methylation in *met1*. Epigenetic remodeling of *met1* recombination is evident at the scale of chromosomes, transposons and individual nucleosomes.

## Discussion

The Arabidopsis meiotic DSB landscape shows both conserved and plant-specific features, compared with SPO11-oligonucleotide maps generated in fungal and mammalian species (Pan et al. 2011; Lange et al. 2016; Fowler et al. 2014). Consistent with the absence of PRDM9 in plants and fungi, Arabidopsis hotspots are more similar to those observed in budding yeast promoters, which are driven by nucleosome occupancy (Pan et al. 2011; Lam and Keeney 2015; Wu and Lichten 1994; Fan and Petes 1996). However, Arabidopsis also shows SPO11-1 hotspots within nucleosome-depleted gene terminators and introns, indicating that varying gene architectures can influence meiotic DSB patterns between species. Interestingly, avian crossover hotspots are also observed at both gene promoters and terminators (Singhal et al. 2015), meaning that recombination hotspots located at gene 3’-ends may be widely conserved.

Consistent with analysis of yeast and mouse SPO11-oligonucleotides, we do not observe a strong relationship between H3K4^me3^ and DSB levels (Tischfield and Keeney 2012; Lange et al. 2016). However, as this modification correlates positively with plant crossover frequency (Choi et al. 2013; Liu et al. 2009; Shilo et al. 2015), it is likely that H3K4^me3^ plays a recombination-promoting role downstream of DSB formation, potentially via tethering repair sites to the chromosome axis, as in budding yeast and mammals (Sommermeyer et al. 2013; Borde et al. 2009; Acquaviva et al. 2013; Imai et al. 2017). No evidence for PRDM9-like proteins exist in plants, which acts to direct recombination hotspots to specific sequence motifs in mammals (Lange et al. 2016; Mihola et al. 2009; Parvanov et al. 2010; Myers et al. 2010; Baudat et al. 2010; Grey et al. 2011; Brick et al. 2012; Grey et al. 2017). However, we observe a strong influence of AT-sequence richness on SPO11-1 levels. As AT-richness excludes nucleosomes (Segal and Widom 2009), we propose that these motifs allow increased SPO11-1 access to DNA, and this underlies their association with elevated crossover frequency (Shilo et al. 2015; Choi et al. 2013; Wijnker et al. 2013).

Several additional differences are notable between the budding yeast and Arabidopsis genomes when comparing their DSB landscapes. First is the possession of point versus regional centromeres (Bloom 2014; Copenhaver et al. 1999; Vincenten et al. 2015), and that Arabidopsis contains a larger and more diverse transposon complement (Buisine et al. 2008; Quadrana et al. 2016; Stuart et al. 2016). Arabidopsis transposons are enriched in pericentromeric heterochromatin and are transcriptionally silenced by DNA methylation (Saze et al. 2003; Kato et al. 2003), which is a chromatin modification not present in budding or fission yeast. Using the *met1* mutant we show that loss of maintenance of CG DNA methylation causes coordinated gain of euchromatic marks, transcription and SPO11-1 DSBs within Arabidopsis centromeric regions. Gain of meiotic DSBs in *met1* was greatest in coldspot transposons, including the EnSpm/CACTA and Gypsy families. Hence, DNA methylation simultaneously silences transcription and initiation of meiotic recombination in specific families of Arabidopsis transposons. This finding is reminiscent of increased SPO11-DSBs detected in specific retrotransposon classes in mouse *dnmt3l* DNA methylation mutants (Zamudio et al. 2015), indicating that epigenetic silencing of transposon recombination is a conserved feature of plant and mammalian genomes.

Despite the expectation that transposons would be recombination-suppressed, in order to avoid genome instability (Sasaki et al. 2010), we show that specific Arabidopsis DNA transposons contain strong meiotic DSB hotspots. These DNA transposons are AT-rich and nucleosome-depleted in wild type and frequently occur in close proximity to genes. As Helitrons and Pogo/Tc1/Mariner transposons display TA and AT dinucleotide insertion site preferences (Guermonprez et al. 2008; Kapitonov and Jurka 2001), this likely contributes to their enrichment in AT-rich gene regulatory regions, where they may further contribute to nucleosome exclusion and enhanced SPO11-1 DSB levels. Higher meiotic recombination initiation may also be responsible for DSB hotspot transposons tending to occur as shorter, non-autonomous fragments. For example, insertions, deletions and rearrangements can result from non-allelic recombination between repeated loci (Sasaki et al. 2010). Together, these data reveal unexpected diversity in the chromatin and recombination landscapes between Arabidopsis transposable element families. As plant genomes vary greatly in the abundance and chromosomal distributions of specific transposon families (Buisine et al. 2008; Choulet et al. 2014; Guermonprez et al. 2008; Kapitonov and Jurka 2001; Quadrana et al. 2016; Stuart et al. 2016; Liu et al. 2009), repetitive elements may contribute to diversity of meiotic recombination patterns between species.

A role for transposons modifying transcription in proximity to genes is well established, consistent with Barbara McClintock’s ‘Controlling Elements’ concept (Slotkin and Martienssen 2007; McClintock 1956). Here we demonstrate that transposons also shape the meiotic DSB and chromatin landscape, within Arabidopsis gene regulatory regions. We propose that nucleosome-depleted SPO11-1 hotspot transposons may provide an adaptive function within plant genomes, by acting as recombination-enhancers. This may be particularly important at the diverse NBS-LRR resistance gene family, which participate in host-pathogen coevolution (Jones and Dangl 2006). Interestingly, these immune loci are also known regions of high genetic and epigenetic divergence between Arabidopsis populations (Alonso-Blanco et al. 2016; Kawakatsu et al. 2016). We propose that hotspot transposons directly contribute to this diversity by recruiting SPO11-1-dependent DSBs during meiosis. Together, our work reveals novel mechanisms whereby mobile genetic elements can influence meiotic recombination, chromatin, diversity and adaptation in their host genomes.

## Methods

### Generation of Arabidopsis *SPO11-1-Myc spo11-1* lines

Six c-Myc (6xMyc) epitopes were translationally fused to a genomic clone of the *SPO11-1* gene in the pPZP211 binary vector, which was transformed into wild type Arabidopsis (Col-0) using *Agrobacterium tumefaciens* strain GV3101, via floral dipping. *SPO11-1-Myc* transformants were crossed with *spo11-1-3* (SALK_146172) heterozygotes to perform fertility complementation tests(Hartung et al. 2007).

### Recombination measurements using fluorescent seed and pollen

Crossover measurements using fluorescent seed or pollen were carried out as described(Ziolkowski et al. 2015; Yelina et al. 2013).

### Immunocytological analysis

Chromosome spreads of Arabidopsis pollen mother cells and immunostaining of ASY1 and SPO11-1-Myc were performed using fresh buds, as described(Armstrong et al. 2002). The following antibodies were used: α-ASY1(Armstrong et al. 2002), (rabbit, 1/500 dilution), α-Myc (9E10, Santa Cruz Biotechnology) (mouse, 1/50 dilution). Microscopy was conducted using a DeltaVision Personal DV microscope (Applied precision/GE Healthcare) equipped with a CDD Coolsnap HQ2 camera (Photometrics). Image capture and analysis was performed using SoftWoRx software version 5.5 (Applied precision/GE Healthcare).

### Immunoprecipitation of SPO11-1-oligonucleotide complexes

Approximately 30 grams of *SPO11-1-Myc spo11-1-3* floral buds were ground to a fine powder in liquid nitrogen and resuspended in 4 volumes of lysis buffer (25 mM HEPES-NaOH pH 7.9, 5 mM EDTA, 1.2% SDS, 1 mM PMSF, 2 mM DTT, 1×Roche Complete Protease Inhibitor Cocktail). The lysis solution was boiled for 20 minutes, followed by rapid chilling on ice. Centrifugation at 4,000*g* for 20 min at 4°C was performed twice and the final supernatant diluted 4-fold by adding 10 mM Tris-HCl pH 8.0, 150 mM NaCl, 1% Triton-X 100. 100 μg of c-Myc Antibody (9E10, sc-40, Santa Cruz) were added to the diluted extract (∼160 ml) in 15 ml tubes and incubated for 8 hours at 4°C with rotation. 1.6 ml of 50% Protein G-Sepharose slurry (71-7083-00, GE Healthcare) was added and incubated overnight at 4°C with rotation. A mock control (no antibody) was performed to validate immunoprecipitation efficiency and specificity at small scale, using western blotting with mouse monoclonal c-Myc antibodies (9E10, sc-40, Santa Cruz) or c-Myc Antibody HRP conjugates (sc-40 HRP, Santa Cruz). Following immunoprecipitation, protein G beads were collected by centrifugation at 500*g* for 1 minute and washed five times with wash buffer (1% Triton X-100, 15 mM Tris-HCl, pH 8.0, 150 mM NaCl, 1 mM EDTA). Immunocomplexes were eluted from the Protein G beads by incubation at 70°C for 15 minutes in 2 volumes of elution buffer (100 mM Tris-Cl, 1 mM CaCl_2_, 10 mM EDTA, 0.5 % SDS). 20 μg/ml of proteinase K was added to the beads and incubated at 50°C for 4 hours with occasional mixing. An equal volume of phenol/chloroform was added to beads, vortexed and centrifuged at 16,000*g* for 10 minutes. The supernatant was transferred to a fresh 1.5 ml tube and phenol/chloroform extraction was repeated. SPO11-1-oligonucleotides were precipitated using 0.1 volume of 3 M sodium acetate pH 5.2,7.5 μg of glycoblue (Ambion AM9515) and an equal volume of isopropanol, followed by incubation at −80°C for 2 hours. SPO11-1-oligonucleotides were collected by centrifugation at 16,000*g* for 45 minutes at 4°C. After two 80% ethanol rinses the pellet was air-dried and resuspended in 30µl of distilled water. 40µl of 2×formamide loading buffer (80% deionized formamide, 10 mM EDTA, pH 8.0, 0.5 mg/ml xylene cyanol FF, 10% saturated bromophenol blue) were added, mixed and incubated at 70°C for 5 minutes. SPO11-1-oligonucleotides and a 20 bp ladder were separated using a 10% TBE-Urea gel (Invitrogen EC6875BOX) and stained by SYBR^®^ Gold Nucleic Acid Gel Stain (Molecular Probes S-11494) for 3 minutes with gentle shaking. The gel region corresponding to 35–50 nt was excised, macerated and soaked in 10 mM Tris (pH 8.0) overnight at 37°C with rotation. The gel fragments were removed by SpinX-centrifuge tube filters (Costar 8163) and the eluate was transferred to fresh 1.5 ml tubes. 0.3 volume of 9 M ammonium acetate, 7.5 μg of glycoblue and 2.5 volumes of 100% ethanol were added, mixed and incubated at −80°C for 2 hours. The size-selected SPO11-1-oligonucleotides were centrifuged at 16,000*g* for 45 minutes as above, rinsed twice by 80% ethanol, air-dried and dissolved in 40 µl of distilled water.

For end-labelling experiments an aliquot (50 μl) of Protein G beads reserved from the immunoprecipitation was washed twice with 1×terminal deoxynucleotidyl transferase (TdT) buffer (50 mM potassium acetate, 20 mM Tris-acetate, 10 mM magnesium acetate, pH 7.9), and incubated with 15 units of TdT (M0315L, NEB), 50 μCi [α-32P]-dCTP triphosphate (5,000 Ci/mmol) and 5 μl of 10×TdT buffer, in a total volume of 50 μl, for 30 minutes at 37°C. The beads were washed three times with wash buffer. SPO11-1-oligonucleotide complexes were eluted by boiling for 3 minutes in 50 μl of 2×Laemmli buffer and separated using a 10% SDS-PAGE gel. The gel was vacuum-dried and radioactivity was detected by exposing to a phosphoimager screen.

### SPO11-1-oligonucleotide library construction

Approximately 1 pmol of purified SPO11-1-oligonucleotides were used for GTP tailing at their 3′-ends. Conditions were used such that between 3 and 5 GMP residues were added per oligonucleotide. A 40 μl reaction was used containing 1×TdT buffer (50 mM potassium acetate, 20 mM Tris-acetate, 10 mM magnesium acetate, pH 7.9), 20 units of TdT (M0315L, NEB), and 2 mM GTP at 37°C for 6 hours. TdT was inactivated by incubating at 75°C for 10 minutes. The G-tailed oligonucleotides were precipitated by incubating with 2.5 volumes of 100% ethanol and 0.3 volumes of 9 M ammonium acetate at −80°C for 2 hours, followed by centrifugation at 16,000*g* for 45 minutes, washing twice with 80% ethanol, air-drying and resuspension in 20 μl of distilled water. G-tailed SPO11-1-oligonucleotides were ligated to a double-stranded DNA adapter in a 40 μl reaction of 1×T4 RNA ligase 2 buffer (50 mM Tris-HCl pH 7.5, 2 mM MgCl_2_, 1 mM DTT, 400 μM ATP), 10 pmol double-stranded 3′ adapter (3’-adapter: top strand, 5′-pTGGAATTCTCGGGTGCCAAGGddC-3′, bottom strand, 5′-AGCCTTGGCACCCGAGAATTCCACCC-3′) (Supplementary Table 14) and 20 units of T4 RNA ligase 2 (dsRNA ligase) (M0239L, NEB) overnight at room temperature. To synthesize complementary strands of SPO11-1-oligonucleotides, 30 μM dNTP and 10 units of Klenow polymerase (NEB) were added to the ligation reaction, incubated at 25°C for 15 minutes, followed by 70°C for 10 minutes. 0.3 volumes of 9 M ammonium acetate, 5 μg of glycoblue and 2.5 volumes of 100% ethanol were added, and DNA precipitated at −80°C for 2 hours, followed by centrifugation at 16,000*g*. The pellet was washed twice with 80% ethanol, air-dried and re-dissolved in 20 µl of water. 30 µl of formamide loading buffer was added, mixed and incubated at 70°C for 5 minutes. The denatured products were separated by electrophoresis using a 10% TBE-Urea gel, and the gel region between 60–80 nt (equivalent to 32–52 nt SPO11-1-oligonucleotides with (rG)3-5 tails and a ligated 23 nucleotide adapter) was excised, macerated and rotated overnight at 37°C overnight in 400 μl of 10 mM Tris-HCl, pH 8.0. The buffer containing dissolved SPO11-1-oligonucleotides was centrifuged through SpinX-centrifuge tube filters. 0.3 volumes of 9 M ammonium acetate, 10 μg of glycoblue, and 2.5 volumes of 100% ethanol were added and DNA was precipitated at −80°C for 2 hours, followed by centrifugation at 16,000*g* for 45 minutes. The pellet was washed twice with 70% ethanol and air-dried. The 3′-ends of gel-purified denatured DNA strands were tailed with GTP by dissolving the dried pellet in a 40 μl tailing reaction containing 1×TnT buffer, 30 units of TdT, and 50 μM GTP, then incubating at 37°C for 6 hours and at 70°C for 10 minutes. The G-tailed products were precipitated with 0.3 volumes of 9 M ammonium acetate, 10 μg of glycoblue, and 2.5 volumes of 100% ethanol at −80°C for 2 hours, followed by centrifugation at 16,000*g* for 45 minutes. After washing with 70% ethanol twice, the air-dried pellet was dissolved in 20 μl of distilled water and incubated in 40 μl of 1×T4 RNA ligase 2 buffer, 10 pmol double-stranded DNA adapter (5′ adapter: top strand, 5′-pATCGTCGGACTGTAGAACTCTGAAddC-3′.bottom strand, 5′-AGTTCAGAGTTCTACAGTCCGACGATCCC-3′) (Supplementary Table 14) and 30 units of T4 RNA ligase2 at room temperature overnight. Finally 30 μM dNTP and 10 units of Klenow polymerase were added and incubated at 25°C for 15 minutes, followed by 70°C for 10 minutes.

A test PCR was performed using a total reaction volume of 20 μl with 1/50 of the final Klenow reaction, 1×FailSafe™ PCR 2×PreMix E (FSP995E, Epicentre), 1μl of Pfu Ultra II Fusion HS DNA Polymerase (Catalog #600672, Agilent) and 1 μM primers RP1 and RPI1. The reaction mixture was divided into two tubes, and PCR performed at 94°C for 20 seconds, followed by 20 cycles of {94°C for 10 seconds; 60°C for 30 seconds; 72°C for 15 seconds}. 5 μl of the PCR products were separated using a 10% TBE gel (EC6275BOX, Invitrogen) with a PCR 20 bp low ladder (P1598, Sigma-Aldrich) and stained with SYBR gold to determine the size and quantity of PCR products. PCRs were then scaled up to a total volume of 400 μl. This mixture was divided into 10 μl aliquots, denatured at 94°C for 10 seconds and amplified for 16 cycles of {94°C for 10 seconds; 60°C for 30 seconds; 72°C for 15 seconds}. PCR products were pooled and precipitated using 0.3 volumes of 9 M ammonium acetate, 7.5 μg of glycoblue and 2.5 volumes of 100% ethanol. The PCR products were separated by electrophoresis using a 10% TBE gel, and the gel area corresponding to 160–180 bp was excised, macerated and soaked in 400 μl of 10 mM Tris, pH 8.0 at 37°C overnight, with mixing. The eluate was spun through a SpinX-centrifuge tube filter and DNA was precipitated using 0.3 volume of 9 M ammonium acetate, 7.5 μg of glycoblue and 2.5 volumes of 100% ethanol. The air-dried DNA pellet was dissolved in 30 μl of 10 mM Tris, pH 8.0. Sequencing was performed using an Illumina NextSeq instrument.

### MNase and H3K4^me3^ chromatin immunoprecipitation sequencing

Micrococcal nuclease digestion and sequencing library construction were performed as reported(Choi et al. 2016). For ChIP two grams of unopened floral buds were ground in liquid nitrogen. Nuclei were isolated and *in vitro* cross-linked in nuclear isolation crosslinking buffer (60 mM Hepes pH 8.0, 1 M sucrose, 5 mM KCl, 5 mM MgCl_2_, 5 mM EDTA, 0.6% Triton X-100, 0.4 mM PMSF, 1 ug pepstatin, 1×protein inhibitor cocktails, 1% formaldehyde) at room temperature for 25 minutes. Glycine was added to a final concentration of 125 mM and incubated for 25 minutes at room temperature with rotation. Cross-linked bud lysate was filtered through one layer of Miracloth and centrifuged at 2,000*g* at 4°C for 20 minutes. The pellet was resuspended in extraction buffer (0.25 M sucrose, 10 mM Tris-HCl pH 8.0, 10 mM MgCl_2_, 1% Triton X-100, 1 mM EDTA, 5 mM β-mercaptoethanol, 0.1 mM PMSF, 1×proteinase inhibitor cocktails) and centrifuged at 2,000*g* at 4°C for 15 minutes. The nuclei pellet was rinsed with 1 ml of TNE buffer (10 mM Tris-HCl pH 8.0, 10 mM NaCl, 1 mM EDTA, 1×proteinase inhibitor cocktails), resuspended and then centrifuging at 2,000*g* at 4°C for 5 minutes. Cross-linked chromatin was digested with 0.05 units of mirococcal nuclease (MNase, NEB M0247S) in reaction buffer (10 mM Tris-HCl, pH 8.0, 10 mM NaCl, 1 mM EDTA, 4 mM CaCl_2_) at 37°C for 15 minutes with vortexing. The reaction was stopped by adding EDTA to a final concentration of 20 mM, vortexing and placing on ice for 10 minutes. One volume of 10 mM Tris-pH 8, 0.2% SDS, 2% Triton X-100, 0.2% sodium deoxycholate, 1×proteinase inhibitor cocktails was added and rotated for 2 hours at 4°C. The reactions were centrifuged at 14,000*g* in a microfuge for 5 minutes at 4°C. The supernatant was used for immunoprecipitation overnight at 4°C using Dynabeads Protein G that were pre-bound to 5 μg H3K4^me3^ antibody (AbCam ab8580). The chromatin immunocomplexes were washed, eluted and reverse-crosslinked. The immunoprecipitates were further purified by phenol/chlorophorm/isoamyl alcohol (24:24:1) extraction, followed by ethanol precipitation and 2% agarose gel separation and gel extraction of ∼145–150 bp DNA. Approximately 10 ng of ChIP-purified DNA was used to generate a library using the TruSeq Prep Kit v2 (Illumina). Libraries were subjected to paired-end sequenced using an Illumina NextSeq instrument.

### RNA-sequencing

Five μg of total RNA from unopened flower buds were extracted using Trizol reagent. To perform rRNA depletion we used the Ribo-Zero magnetic kit (MRZPL116). Fifty ng of rRNA-depleted RNA were used for RNA-seq library construction using the ScriptSeq v2 RNA-seq Library Preparation Kit (SSV21124). The library was amplified using 12 PCR cycles and indexed using ScriptSeq Index PCR Primers (RSBC10948) and FailSafeTM PCR Enzyme Mix (FSE51100). Sequencing was performed on a HiSeq instrument. RNA-seq data were analyzed using RSem.

### Bioinformatics analysis of SPO11-1-oligonucleotides, ChIP-seq and MNase-seq data

For SPO11-1-oligonucleotide data FASTQ files were trimmed for 3′-adapter sequences using the FASTX-Toolkit function fastx_clipper (http://hannonlab.cshl.edu/fastx_toolkit/). For the wild type libraries RPI1 and RPI3 5 bp were trimmed from the read 5′-ends, while for other libraries 10 bp were cropped, due to longer adapter sequences. Trimmed reads were aligned to the TAIR10 reference sequence using bowtie2 with the following settings: --very-sensitive -p 4 -k 10. Aligned reads were filtered to have 2 or fewer mismatches. Reads with the SAM optional field “XS:i” were dropped to obtain unique alignments. Reads with multiple valid alignments were filtered for MAPQ scores of 10 or higher, and the highest value alignment kept. In the event that a read had multiple alignments with equal MAPQ scores, one was randomly chosen. Unique and multiply aligning reads were then deduplicated using SAMtools. BAM files for uniquely and multiply aligning reads were combined. For MNase-seq and ChIP-seq data paired-end FASTQ files were directly aligned to the TAIR10 reference sequence using bowtie2 with the following settings -very-sensitive -no-discordant -no-mixed -p 4 -k 10. To obtain uniquely aligning reads, reads with the SAM optional field “XS:i” and MAPQ scores of less than 42 were dropped. To ensure reads were kept in proper pairs, a Python script was applied. Reads with multiple valid alignments were filtered for those with MAPQ scores of 10 or higher and the highest value alignments kept. Multiply aligning reads were treated as for SPO11-1-oligonucleotides. Unique and multiply aligning reads were then deduplicated using SAMtools, combined and used for downstream analysis. Coverage values from these reads were calculated using Rsamtools and normalized by the sum of coverage per library. Analysis of these data in relation to features including TAIR10 representative gene TSS and TTS and transposons(Buisine et al. 2008), was performed as previously described (Choi et al. 2013). For hexile analysis normalized values of SPO11-1-oligonucleotides were calculated in windows −500 bp upstream of TSS for promoters or +500 bp downstream of TTS for terminators. These regions were also measured for nucleosome occupancy and AT-rich motif matches. These were compared with H3K4^me3^ and CTT motif matches in the 500 bp downstream of TSS and upstream of TTS. To test the extent of SPO11-1-oligonucleotide hexile overlap with crossovers, we used a set of 2,499 crossovers mapped using genotyping-by-sequencing in Col×Ler F_2_ individuals(Choi et al. 2016; Yelina et al. 2015). SPO11-1 levels were calculated within each SNP interval used to detect crossovers. Intervals were also classified according to their overlap with genomic annotation and position along chromosomes in 2 megabase bands. Data were modeled with the glm function in R, using the binomial family and a logistic link function.

### Data Access

The FASTQ files associated with the genomic datasets described here have been uploaded to the ArrayExpress repositories, and can be accessed using the provided usernames and passwords.

SPO11-1-oligonucleotides: <u>https://www.ebi.ac.uk/arrayexpress/experiments/E-MTAB-5041/</u>

Username: Reviewer_E-MTAB-5041 Password: MKE8bvew

Nucleosome MNase-seq: <u>https://www.ebi.ac.uk/arrayexpress/experiments/E-MTAB-5042/</u>

Username: Reviewer_E-MTAB-5042 Password: 4c0zvhju

H3K4^me3^ ChIP-seq: <u>https://www.ebi.ac.uk/arrayexpress/experiments/E-MTAB-5048/</u>

Username: Reviewer_E-MTAB-5048 Password: 4c0zvhju

RNA-seq: <u>https://www.ebi.ac.uk/arrayexpress/experiments/E-MTAB-5417/</u>

Username: Reviewer_E-MTAB-5417 Password: 2rX8I48v

## Acknowledgements

Research was supported by a Royal Society University Research Fellowship, the Gatsby Charitable Foundation grant GAT2962, BBSRC grant BB/N007557/1, National Natural Science Foundation of China grant 61403318, Next-Generation BioGreen Program (SSAC grant PJ01137901 RDA Korea) and an EMBO long-term postdoctoral fellowship (ALT 807-2009).

## Author Contributions

KC, CL, CJU, HS, PAZ, NEY, RAM and IRH contributed to design of the study. KC, CL, CJU and HS performed experiments. KC, XZ, CL, CJU, TJH, HS, AJT, RAM and IRH analyzed the data. KC, XZ, CL CJU, TJH, HS, AJT, PAZ, NEY, RAM and IRH wrote the manuscript.

## Supplemental Material

Supplemental Figures S1–S6

Supplemental Tables S1–S14

